# Effects of CEPA and 1-MCP on flower bud differentiation of apple cv. ‘Nagafu No.2’ grafted on different rootstocks

**DOI:** 10.1101/281303

**Authors:** Wenfang Li, Baihong Chen, Juan Mao, Xinwen Li, Jing Su, Mohammed Mujitaba Dawuda, Zonghuan Ma, Cunwu Zuo, Zeshan An

## Abstract

The apple (*Malus domestica* Borkh.) has a relatively long juvenile period which prevent the fruit breeding. The understanding of the flowering system is important to improve breeding efficiency in the apple. In this context, 2-year-old “Fuji” apple cv. “Nagafu No.2” trees that were grafted on dwarf self-rooted rootstock M.26, vigorous rootstock *M. sieversii* and interstock M.26/*M. sieversii*, respectively. Spraying with clean water (as controls), 800 mg·L^−1^ 2-Chloroethylphosphonic acid (CEPA) and 2 μL·L^−1^ 1-methylcyclopropene (1-MCP). The results showed that CEPA significantly repressed the vegetative growth attributed to the increase of the ABA and ZT synthesis, and the decrease of IAA synthesis in leaves and buds. However, there was no significant difference or significant inverse effect between 1-MCP and control. Furthermore, CEPA promoted flower formation, increased the flowering rate and advanced the blossom period for 2 days compared with the control, which accompanied by the accumulation of soluble sugar, glucose and sucrose, and the increase of α-amylase (α-AMY) and sucrose phosphate synthase (SPS) activities, and the decrease of the starch contents and sucrose synthase (SS) activities in leaves and buds. However, the blossom period was delayed for 2 days after spraying with 1-MCP. Finally, the expression of *TFL1* was significantly repressed while the *AP1* was significantly promoted in buds from M.26 and M.26/*M. sieversii* after spraying with CEPA, while the effect was not significant from *M. sieversii.* However, the expression levels of *TFL1* and *AP1* were not significantly different from the control after the application of 1-MCP. In spite of this, CEPA was more susceptible to easy-flowering M26, followed by M26/*M. sieversii*, and still less susceptible to difficult-flowering rootstock *M. sieversii*.

**Abbreviations:**

1-MCP: 1-methylcyclopropene
α-amylase: (α-AMY)
ABA: abscisic acid
CEPA: 2-Chloroethylphosphonic acid
CTK: cytokinins
ETH: ethylene
GA: gibberellin
SPS: sucrose phosphate synthase
SS: sucrose synthase
ZT: zeatin.

## Introduction

The cultivated apple (*Malus domestica* Borkh.) is one of the commercially important fruit crops in the world and has a relatively long juvenile growth period, which disturbs early cropping and fruit breeding (Dennis *et al*., 2003; Mimida *et al*., 2012). Moreover, it takes from 14 to 16 months for a reproductive period between the flower initiation and the mature fruit in the apple (Mimida *et al*., 2011). The flower initiation occurred in the early summer of the previous year affects directly the yield of apple crops in the autumn of the following year. Apple cv. ‘Nagafu No.2’ is a good case in point. Therefore, the understanding of the flowering system is important to improve breeding efficiency and maintain a steady harvest in the apple.

Rootstock effects on apple scions with different growth habits and gene expression patterns (Jensen *et al*., 2003; Tworkoski and Miller, 2007). Interstock bridge grafting of mature apple trees maybe a viable method of reducing vegetative growth with increased reproductive growth and improving yield (Samad *et al*., 1998). Dwarfing root-stock would decrease branch angles, inhibit stem elongation more inupright than in wide angled trees which makes it more convenient to manage and increase productivity (Tworkoski and Miller, 2007). However, interstock bridge grafting should be selected in the areas without irrigation conditions, and both the interstock and the self-rooted rootstock can be used in areas with irrigation conditions (Zhang *et al*., 2016). The common rootstock used in dwarfing cultivation is M, MM, P, B and and SH (Tworkoski and Miller, 2007; Gjamovski and Kiprijanovski, 2011). However, ‘Fuji’ apple is still more difficult to be flowering than other apple cultivars even if grafted on the dwarfing root-stock.

The level of sugar metabolism in the organs of the fruit trees has a great influence on the flower bud differentiation. Studies have shown that the metabolism, transport and accumulation of carbohydrates are significantly related to flower bud differentiation in fruit trees (Green *et al*., 1993; Ito *et al*., 2002; Eshghi *et al*., 2007). Furthermore, flower bud differentiation is also influenced by the type and content of plant endogenous hormones in fruit trees (Porri *et al*., 2012, Li *et al*., 2016). Phytohormones play important roles in flower bud differentiation (Kong *et al*., 2009; Li *et al*., 2016). Moreover, most of the genes regulating flower development have been found, mainly represented by *FLOWERING LOCUST* (*FT*), *TERMINAL FLOWER1* (*TFL1*), *LEAFY* (*LFY*), *APETALA1* (*AP1*), *CONSTANS* (*CO*) and *EMBRYONIC FLOWER* (*EMF*) (Moon *et al*., 2003; Wang *et al*., 2007; Kotoda *et al*., 2010; Sadao and Masato, 2012; Haberman *et al*., 2016; Patil *et al*., 2017). In apple, the counterparts of the floral genes including *MdFT, MdTFL1*, and *MdAP1*, had been isolated and characterized (Hättasch *et al*., 2008; Flachowsky *et al*., 2009; Kotoda *et al*., 2010; Zhang *et al*., 2016).

The temperate deciduous tree apple, flowers autonomously, with floral initiation dependent on aspects of vegetative development in the growing season before anthesis (Lee and Safe, 2008). Delay in flowering is one of the major limitations in the production of “Fuji” apple (Fan *et al*., 2017). Ethephon (2-Chloroethylphosphonic acid, CEPA) is an ethylene (ETH) yielding chemical, when applied directly to plants, can elicit a response characteristic of ETH treatment (Yang, 1969; Cools *et al*., 2011). 1-Methylcyclopropene (1-MCP) is an anti-ETH compound due to its ability to block the ETH binding sites in plant cells (Prange and DeLong, 2003; Prange *et al*., 2005). Hormones such as indole-3-acetic acid (IAA), cytokinins (CTK), abscisic acid (ABA) and gibberellic acid (GA), play important roles in the flower bud induction of fruit trees (Dieleman *et al*., 1998; Porri *et al*. 2012, Li *et al*. 2016). Nevertheless, the effect of exogenous ETH and its inhibitor 1-MCP on flower bud differentiation in fruit trees was rarely reported. In the present research, 2-year-old “Fuji” apple cv. “Nagafu No.2” was grafted on dwarf self-rooted rootstock M.26, vigorous rootstock *M. sieversii* and interstock M.26/*M. sieversii*, respectively. CEPA and 1-MCP were used to compare the physiological and biochemical differences among scion-stock combinations. The effect of CEPA and 1-MCP on the flower bud differentiation of apple from scion-stock combinations interactions with vegetative growth, carbohydrates, and plant endogenous hormones, are discussed, in order to provide a theoretical basis for solving the problems of late flowering and explore the regulation mechanism of plant growth regulators on flower bud differentiation in fruit trees.

## Materials and methods

### Plant materials, growth conditions and treatments

Experiments were conducted in 2016 and 2017, using 2-year-old Fuji apple cv. ‘Nagafu No.2’ trees that were grafted on dwarf self-rooted rootstock M.26, vigorous rootstock *M. sieversii* and interstock M.26/*M. sieversii* as research materials. Trees were planted in the Apple Demonstration Nursery in Qingcheng county, Gansu Province of China (36°14N, 107°88E). The plants were at 1.5 × 4.0 m row and plant spacing for both dwarf self-rooted rootstock and interstock, 3.0 × 5.0 m for vigorous rootstock which were managed using standard horticultural practices. Spraying clean water on July 1^st^, 2016 until dripping on the branches on the whole trees of scion-stock combinations including Nagafu No.2/M.26, Nagafu No.2/*M. sieversii* and Nagafu No.2/M.26/*M. sieversii* as controls, were named as CK1, CK2 and CK3, respectively. Additionally, spraying scion-stock combinations with 800 mg·L^−1^ CEPA and 2 μL·L^−1^ 1-MCP as treatments, were named as A1, A2, A3 and B1, B2, B3, respectively. Furthermore, the entire tree with a black plastic bag for 24 hours after spraying. The flower bud morphology was measured twice at 16 days (17^th^ July, 2016) and 31 days (2^nd^ August, 2016) after spraying treatments. The physiological and biochemical indicators including sugar content, enzyme activity and hormone content were measured three times, at 1 day (2^nd^ July, 2016), 16 days and 31 days after spraying treatments. Samples for RNA extraction were collected once at 16 days after spraying treatments. Samples (buds and their surrounding leaves) for measuring physiological and biochemical indicators or RNA extraction were immediately frozen in liquid nitrogen and stored at −80 °C. Each treatment was subdivided into three subgroups consisting of 30 trees each to allow for 3 replications. Flowering rates were calculated during the blossom period, 298 days after spraying (April 28, 2017).

### Measurement of growth indicators

The length of terminal shoots (above the interface) and new shoots was measured by tape measure. The stem diameter (10 cm above the interface) was measured with a vernier caliper. Dry and fresh weight of leaves was measured by electronic balance. The increment of these growth indicators comes from 31 days to 1 day.

### Scanning electron microscopy

Fifteen fresh flower buds which were uniform in size and vigor were collected. The samples were fixed and stored in serum bottles containing 10 mL of 2.5% glutaraldehyde fixative and placed in a vacuum pump for 30 min. Twelve flower buds were selected and rinsed three times (10 min each) in 0.1 mol·L^−1^ phosphate buffer and were kept in 50% ethanol during dissection to prevent desiccation. Later, the samples were dehydrated in ethanol series (one time 30%, 50%, 70%, 80% and 90% for 30 min and then three times in 100% for 15 min). And then dehydrated with tert-butyl alcohol instead of ethanol, and the dehydration process cannot be reversed. After 3 h of air-drying, peeling flower bud, part of calyx and hair flake under stereo microscop. These samples were glued to the stage one by one with conductive adhesive, which were coated with palladium-gold in a sputter coater (S150B; Edwards) for about 1 min, were examined by scanning electron microscope (S-3400N; Hitachi) at an acceleration voltage of 10 kV.

### Starch and sugars content measurements

Starch content was measured by acid hydrolysis method given by McCreddy *et al*. (1950). To the residue, 5mL of distilled water and 6.5 mL of 52% perchloric acid was added to extract the starch by placing the samples at 0 °C for 20 min. The mixture was centrifuged and retained the extract. The process was repeated 3-4 times using fresh perchloric acid and diluted to final volume 100 mL. To 0.5 mL of diluted extract, 4.5 mL of distilled water was added followed by addition of 10 mL of cold anthrone sulfuric acid reagent in ice bath. The sample mixture was heated at 100 °C for 8 min, and cooled rapidly to room temperature. The absorbance was measured at 630 nm. The final content of starch was calculated from a standard curve plotted with known concentration of glucose.

Soluble sugars were determined Spectrophotometrically by anthrone reagent using glucose as standard at 625 nm (Dubois *et al*., 1956). Approximately 0.2 g of frozen bud and leaf samples were used to measure the content of glucose and sucrose by kit from Suzhou Comin Biotechnology Co., Ltd (Suzhou, China) according to the instructions. They were finally assayed from the obtained light absorption value at 480 nm.

### Enzyme activity measurements

Activity of α-amylase (α-AMY) was assayed by the method of Shuster and Gifford (1962). Approximately 0.5 g of frozen Frozen bud and leaf samples were homogenized in ice cold extraction buffer (0.1 M phosphate buffer pH 7.0), centrifuged at 10,000 rpm (4 °C) and the supernatant treated as enzyme extract. One mL of starch substrate was added to 0.5 mL of enzyme extract. 0.2 mL of aliquot was removed from this and added 3 mL of KI. The absorbance was recorded at 620 nm. Then, the reaction mixture left was incubated at 25 °C, and then removed the aliquot and repeated the color developing process (violet blue) after every 30 min. Blank was run simultaneously without having substrate. In control the enzyme extract was substituted with 0.5 mL of distilled water.

Approximately 0.1 g of frozen bud and leaf samples were used to measure sucrose synthase (SS) and sucrose phosphate synthase (SPS) activities by enzyme activity kit from Suzhou Comin Biotechnology Co., Ltd (Suzhou, China). The 1 mL extract was added for grinding, centrifuged at 8,000 *g* for 10 min (4 °C). The supernatant was placed on the ice to be measured, and then follow the instructions. SS and SPS activities were assayed from the obtained light absorption value at 480 nm.

### Hormone measurements

Approximately 0.5 g of frozen bud and leaf samples were used to measure IAA, ABA and zeatin (ZT). Each sample was then combined with 10 mL of 80% chromatographic pure methanol (preparation with DNase/RNase-free double-distilled water). Each sample was washed thrice with solvent, transferred into a test tube, and stored in a refrigerator at 4 °C overnight in the dark. Then, the samples were centrifuged for 20 min under refrigerated conditions at 4 °C. Supernatant fluid was transferred into a new centrifuge tube. The extract was concentrated, and the methanol was volatilized under 40 °C by rotary evaporation to obtain 2 mL of concentrate. The evaporation bottle wall was then washed continuously with 50% methanol, and the volume was raised to 10 mL with 50% chromatographic pure methanol. The fluid for testing was filtered through 0.22 μm organic membrane. This fluid was then transferred to a 2 mL centrifuge tube and placed in an ice box. The hormone contents from the treatments were identified in the Instrumental Researches and Analysis Center of Gansu Agricultural University.

The determination method was performed with different concentrations of IAA, ABA, and ZT, standard samples, which were used to construct a standard curve. The standard samples were purchased from Sigma Company, and the external standard curve and quantitative methods were performed for the measurements. The type of LC–MS apparatus used was the Agilent 1100 series (Agilent Technologies, Waldbronn, United States). The detector was vwd, and the chromatographic column was Extend-C18 (4.6 mm × 250 mm, 5 μm). The mobile phase was chromatographic methanol and 0.6% iced acetic acid previously subjected to ultrasonication (0 min, methanol:acetic acid = 40:60; 11.9 min, methanol:acetic acid = 40:60; 12 min, methanol:acetic acid = 50:50). The flow velocity was 1.0 mL/min, the wavelength was 254 nm, and the column temperature was 25 °C.

### RNA extraction and cDNA synthesis

The RNA plant reagent (Real-Times Biotechnology, Beijing, China) was used to extract floral bud total RNA. RNase-free DNase I (Takara, Dalian, China) was used to purify RNA. During RNA purification, DNase digestions were performed three times. 1% (w/v) agarose gel analysis was used to assess RNA quality. Quantification of RNA was performed using the absorbance at 260 nm. After that, quantified RNA was reverse transcribed into cDNA using the Super-Script First-Strand Synthesis system (Invitrogen, Carlsbad, CA).

### Gene expression analysis by qRT-PCR

Genes including terminal flower 1-like (*TFL1*) and apetala 1-like (*AP1*) were chosen for validation using qRT-PCR. *MdGADPH* was used as internal reference. Primers for qRT-PCR, which were designed with Primer 5.0 software, are shown in Table S1. qRT-PCR reactions were analyzed in ABI StepOne ^TM^ Plus Real-Time PCR System with SYBR Green PCR Master Mix (Takara, Dalian, China), and amplified with 1 μL of cDNA template, 10 μL of 2 × SYBR Green Master Mix, and 1 μL of each primer, to a final volume of 20 μL by adding water. Amplification program consisted of one cycle of 95 °C for 30 s, 40 cycles of 95 °C for 5 s, and 60 °C for 34 s, followed by one cycle of 95 °C for 15 s, 60 °C for 60 s, and 95 °C for 15 s. Fluorescent products were detected in the last step of each cycle. Melting curve analysis was performed at the end of 40 cycles to ensure proper amplification of target fragments. All qRT-PCR for each gene was performed in three biological replicates, with three technical repeats per experiment. Relative gene expression was normalized by comparing with CK expression and analyzed using comparative 2^−ΔΔC^T Method (Livak and Schmittgen, 2001). The results of gene expression were visualized by histogram and heatmap.

### Statistical analysis

Data on growth, physiological parameters and expression levels of the flowering genes in the buds and leaves of ‘Nagafu No.2’ were analyzed by one-way analyses of variance and means separated by Duncan multiple range test at *P* < 0.05 using SPSS software, version 21.0 (SPSS, Chicago, IL, USA). Figures were prepared using Origin 9.0 (Microcal Software Inc., Northampton, MA, USA).

## Results

### Effects of CEPA and 1-MCP on the growth parameters of apple from scion-stock combinations

After 31 days of spraying CEPA, the increment of terminal shoot length and new shoot length from A1, A2 and A3 was significantly lower than those of CK1, CK2 and CK3, respectively (Table 1). After 31 days of spraying 1-MCP, the increment of terminal shoot length and new shoot length of B1, B2 and B3 were not significantly different from those of CK1, CK2 and CK3, respectively, except that the increment of terminal shoot length of B1 was significantly higher than that of CK1. However, there is no significant difference in the increment of new shoot diameter between the different treatments from the same rootstock. Furthermore, the overall trend of the three growth parameters from the three different scion-stock combinations under the same treatment was the vigorous rootstock *M. sieversii*, followed by the interstock M.26/*M. sieversii*, and finally the dwarf self-rooted rootstock M.26.

**Table 1.** Effects of CEPA and 1-MCP on growth parameters of apple cv. ‘Nagafu No.2’.

|  | Increment of terminal shoot length (cm) | Increment of diameter of new shoot (mm) | Increment of new shoot length (cm) |
| --- | --- | --- | --- |
| A1 | 16±0.65e | 2.93±0.19b | 6.02±0.24d |
| B1 | 24±0.83c | 2.91±0.15b | 8.14±0.78bc |
| CK1 | 22±0.69d | 2.87±0.07b | 7.80±0.49c |
| A2 | 22±0.43d | 3.92±0.33a | 8.72±0.61bc |
| B2 | 29±0.97a | 3.83±0.31a | 13.36±0.96a |
| CK2 | 28±0.86a | 3.85±0.13a | 11.83±1.18a |
| A3 | 21±0.93d | 3.01±0.35b | 6.46±0.63d |
| B3 | 25±0.87b | 3.84±0.30a | 9.08±0.37b |
| CK3 | 25±0.11b | 3.53±0.24ab | 8.89±0.44b |

Note: Different small letters indicate significant difference at 0.05 levels.

### Effects of CEPA and 1-MCP on the bud morphology of apple from scion-stock combinations

On the 16th day after spraying CEPA, the central protruding part of the growth point from A1, A2 and A3 was significantly changed compared with CK1, CK2 and CK3, respectively (Fig. 1A). However, the growth points of B1, B2 and B3 were not significantly different from those of CK1, CK2 and CK3, respectively.

**Fig. 1.**
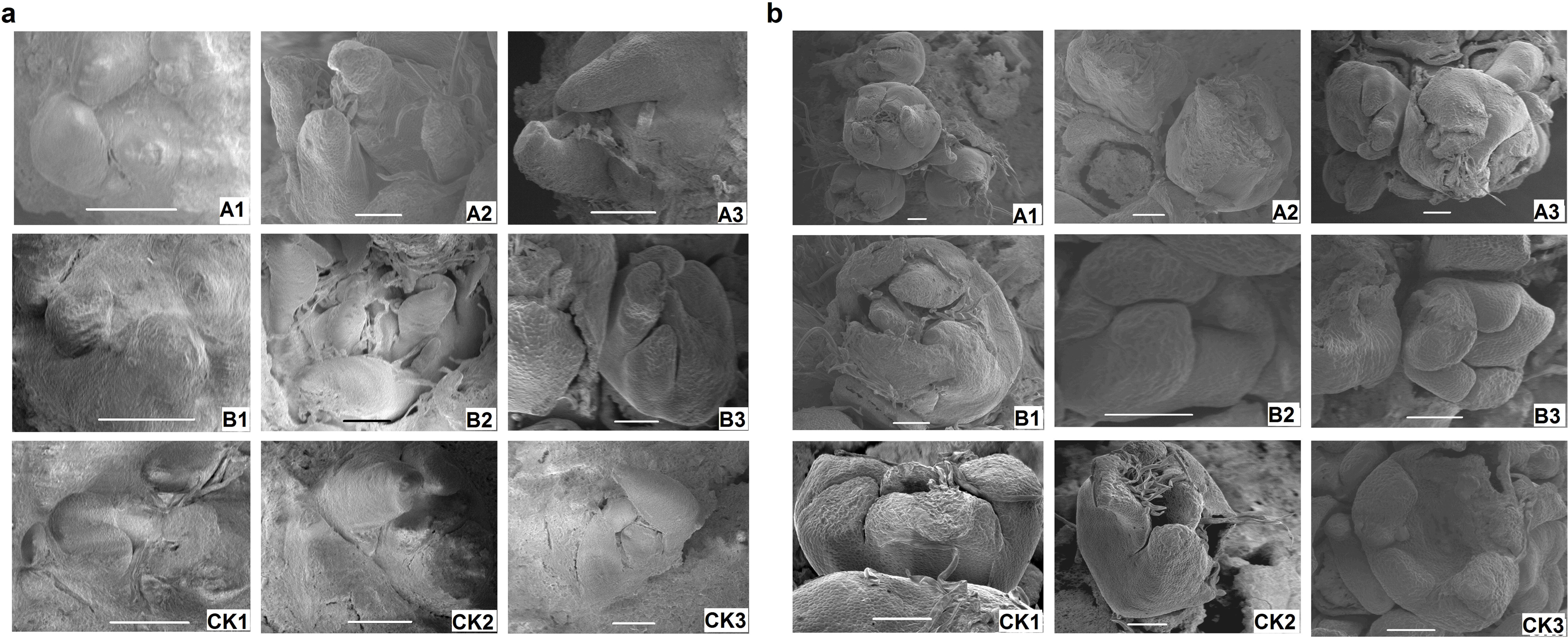
Changes of flower bud morphology from 3 scion-stock combinations including Nagafu No.2/M.26, Nagafu No.2/*M. sieversii* and Nagafu No.2/M.26/*M. sieversii* after exposure to 800 mg·L^−1^ 2-Chloroethylphosphonic acid (CEPA) and 2 μL·L^−1^ its inhibitor of 1-methylcyclopropene (1-MCP) after 16 days (a) and 31 days (b) under scanning electron microscope. Bars=100 μm.

After 31 days of spraying with CEPA, A1, A2 and A3 were differentiated into stamen primordia within two adjacent petal primordia, forming 5 protrusions, that is, the pistil primordium (Fig. 1B). Meanwhile, the development of the primitive lateral flower bud begins with the differentiation of 2 stipules primordia from both sides of the original lateral flower bud, and then 5 sepal primordia were gradually carried out, forming the lateral flower bud in the same order as the central flower bud. There were significant differences in flower bud morphological changes compared to CK1, CK2 and CK3, respectively. In addition, after spraying with 1-MCP, the lateral flowering primordium of B2 appeared, and entered the late stage of flower bud differentiation, which was significantly different from the CK2. The growth cone of CK2 was further uplifted and the sepal primordium was formed in the periphery. The growth cone of B3 was further uplift, formed in the periphery of 5 sepals primordia, and the difference was significant with that of CK3.

### Effects of CEPA and 1-MCP on flowering rate of apple from scion-stock combinations

According to the survey on flowering of ‘Nagafu No.2’ in 2017, the average flowering rate per tree of A1 and A3 were significantly higher than CK1 and CK3, respectively (Fig. 2). Whereas the flowering rates of B1 and B3 were significantly less than CK1 and CK3, respectively. However, the flowering rates of A2 was close to 0, both B2 and CK2 were 0. CK1 and CK3 reached the full flowering stage on May 1^st^. Both A1 and A3 reached full flowering on April 28^th^, and all the calluses were fully expanded. Blossom of B1 and B3 was delayed, and flower buds did not fully expand on April 28^th^ until May 3^rd^. In addition, the overall order of flowering rate of the three different scion-stock combinations was the dwarf self-rooted rootstock M.26, followed by the interstock M.26/*M. sieversii*, and finally the vigorous rootstock *M. sieversii*.

**Fig. 2.**
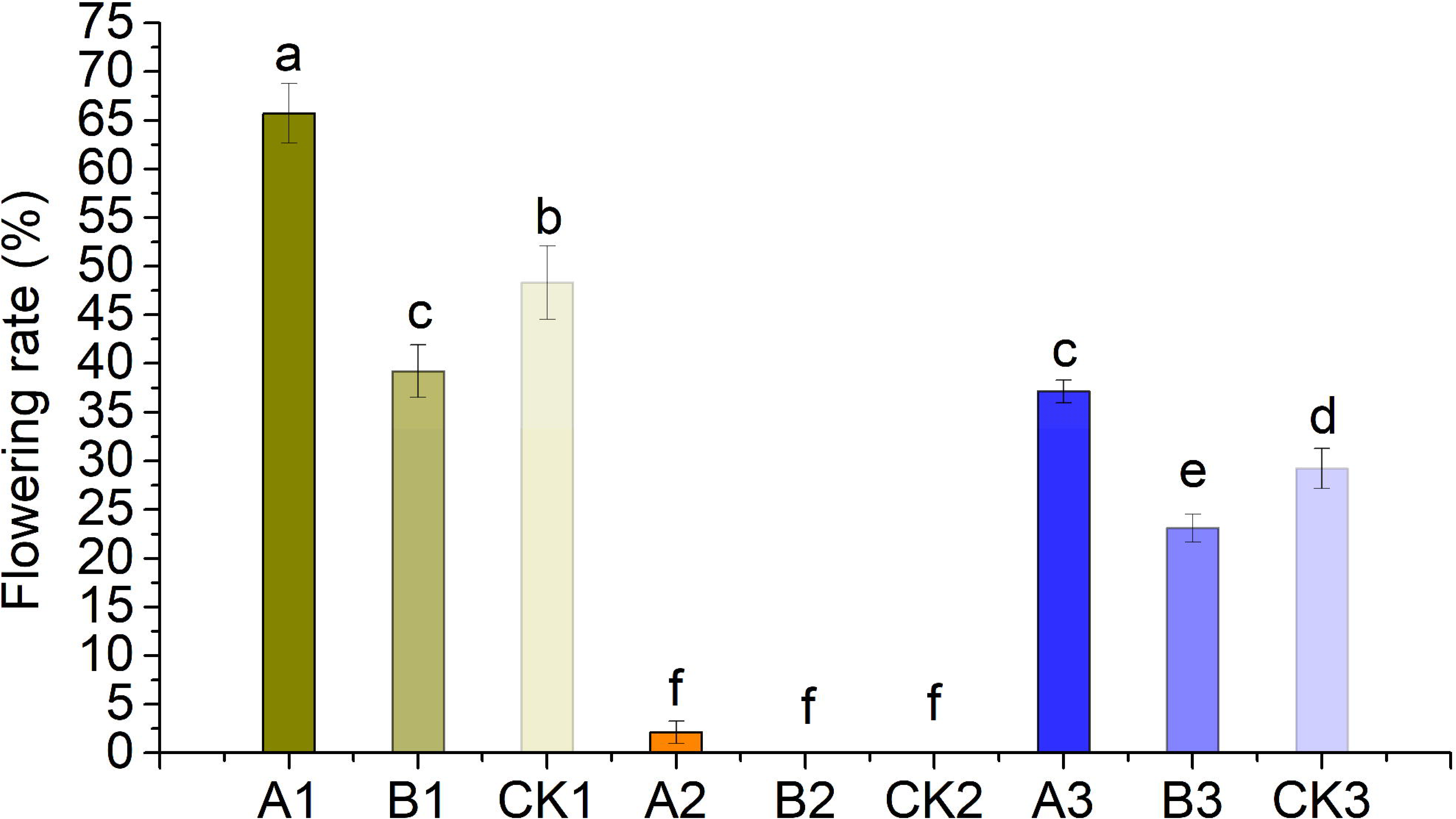
Effects of 800 mg·L^−1^ 2-Chloroethylphosphonic acid (CEPA) and 2 μL·L^−1^ its inhibitor of 1-methylcyclopropene (1-MCP) on flowering rate of ‘Nagafu No.2’. Bars indicate SE. Different small letters within the figures indicate significant difference at 0.05 levels.

### Effects of CEPA and 1-MCP on starch and sugars content in buds and leaves of apple from scion-stock combinations

The starch contents in buds and leaves of all treatments were decreased slightly at 31 days after spraying compared with 16 days. There was no significant difference in starch content from buds and leaves a day after spraying treatments from the same scion-stock combination (Fig. 3A, B). The starch contents were significantly lower in A1, A2 and A3 than those of CK1, CK2 and CK3 in buds and leaves from the 16 and 31 days after spraying, respectively. On the contrary, the starch contents in buds and leaves of B1, B2 and B3 were significantly higher than those of CK1, CK2 and CK3 or no significant difference, respectively. The contents of soluble sugar, glucose and sucrose in buds and leaves increased first and then decreased slightly from day 1 to 31 after spraying treatments, and there was no significant difference a day after spraying treatments from the same scion-stock combination (Fig. 3C–H). Furthermore, the contents of soluble sugar, glucose and sucrose were significantly higher in buds and leaves of A1, A2 and A3 than those of CK1, CK2 and CK3 from 16 and 31 days after spraying, respectively. However, they were significantly lower in B1, B2, B3 than in CK1, CK2 and CK3 or no significant difference, respectively. Additionally, the overall trend of the contents of soluble sugar, glucose and sucrose in buds and leaves of the three scion-stock combinations under the same treatment was dwarf self-rooted rootstock M.26, followed by the interstock M.26/*M. sieversii*, and finally the vigorous rootstock *M. sieversii*. On the contrary, the starch content showed the opposite trend.

**Fig. 3.**
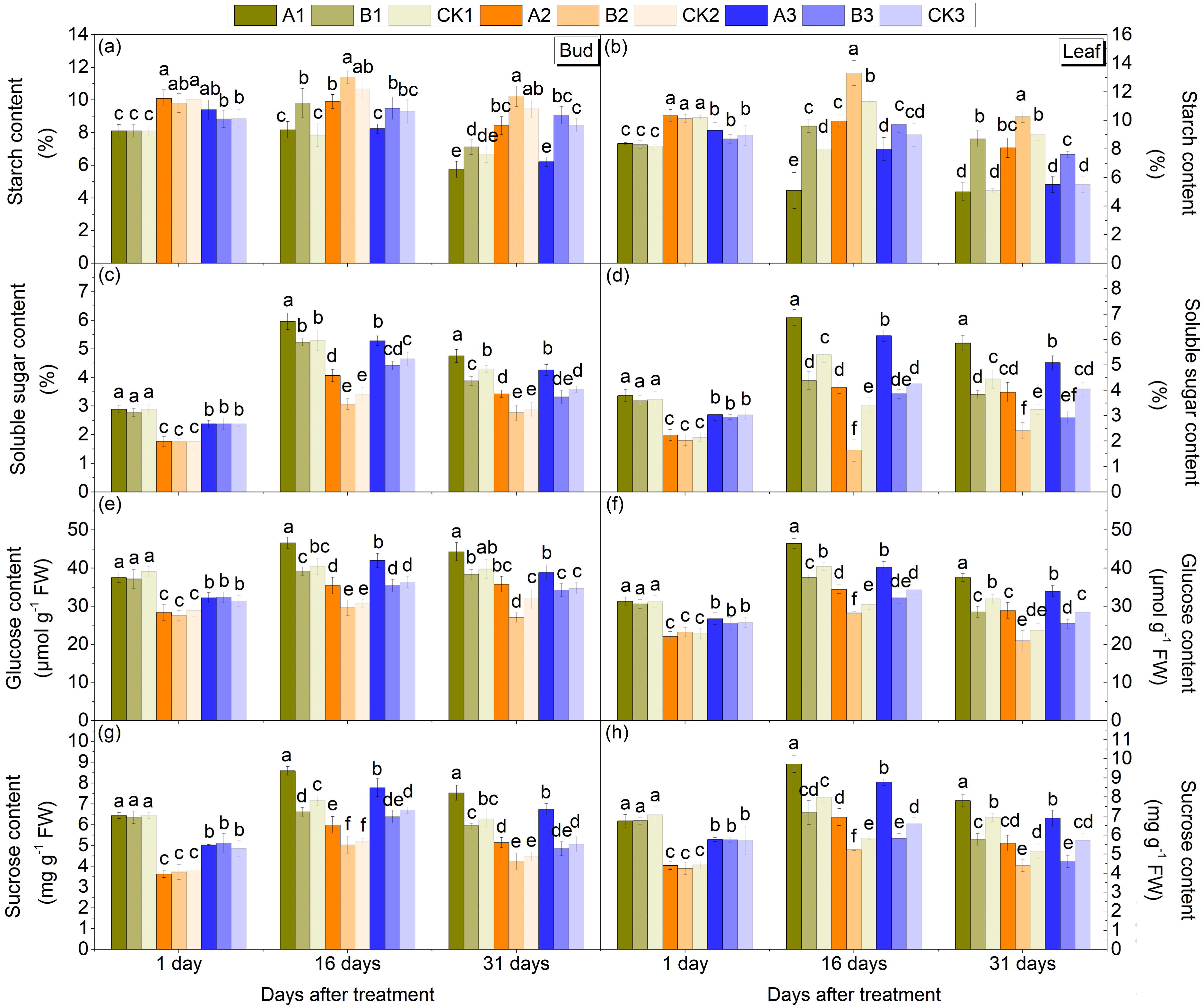
Changes of starch (a and b), soluble sugar (c and d), glucose (e and f) and sucrose (g and h) contents in buds and leaves of 3 scion-stock combinations including Nagafu No.2/M.26, Nagafu No.2/*M. sieversii* and Nagafu No.2/M.26/*M. sieversii* after exposure to 800 mg·L^−1^ 2-Chloroethylphosphonic acid (CEPA) and 2 μL·L^−1^ its inhibitor of 1-methylcyclopropene (1-MCP) after 1, 16 and 31 days. Bars indicate SE. Different small letters within the figures indicate significant difference at 0.05 levels.

### Effects of CEPA and 1-MCP on activities of enzymes related to sugar metabolism in buds and leaves of apple from scion-stock combinations

There was no significant difference in activities of α-AMY from buds and leaves a day after spraying treatments from the same scion-stock combination. In addition, the activities of α-AMY in buds and leaves of A1, A2 and A3 from 16 and 31 days were significantly higher than those of CK1, CK2 and CK3, respectively (Fig. 4A, B). However, they were significantly lower in buds and leaves of B1, B2 and B3 than those of CK1, CK2, CK3 or no significant difference, respectively. The activities of SS also were not significantly different in buds and leaves a day after spraying treatments from the same scion-stock combination, but significantly lower in A1, A2 and A3 from 16 and 31 days after spraying than those in CK1, CK2 and CK3, respectively (Fig. 4C, D). However, B1, B2 and B3 were significantly higher than those of CK1, CK2 and CK3 or no significant difference, respectively. Remarkably, SPS activity showed the opposite trend with SS activity (Fig. 4E, F). The activities of α-AMY, SS and SPS in buds and leaves of all treatments increased from day 1 to 31 days. Further, the overall trend of α-AMY, SS and SPS activity in buds and leaves of the three scion-stock combinations under the same treatment also was dwarf self-rooted rootstock M.26, followed by the interstock M.26/*M. sieversii*, and finally the vigorous rootstock *M. sieversii*.

**Fig. 4.**
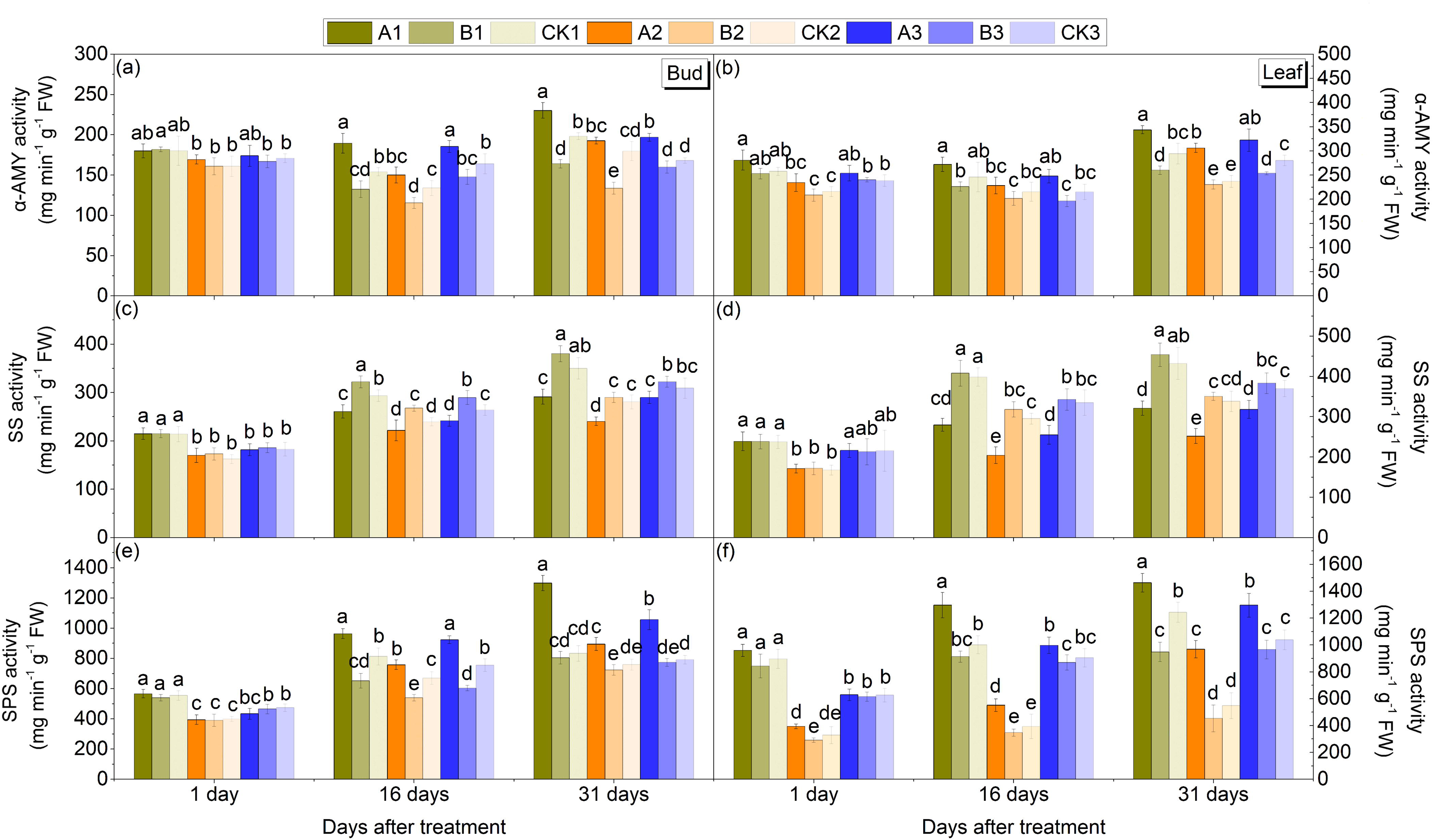
Changes of α-amylase (α-AMY) (a and b), sucrose synthase (SS) (c and d) and sucrose phosphate synthetase (SPS) (e and f) activity in buds and leaves of 3 scion-stock combinations including Nagafu No.2/M.26, Nagafu No.2/*M. sieversii* and Nagafu No.2/M.26/*M. sieversii* after exposure to 800 mg·L^−1^ 2-Chloroethylphosphonic acid (CEPA) and 2 μL·L^−1^ its inhibitor of 1-methylcyclopropene (1-MCP) after 1, 16 and 31 days. Bars indicate SE. Different small letters within the figures indicate significant difference at 0.05 levels.

### Effects of CEPA and 1-MCP on hormones content in buds and leaves of apple from scion-stock combinations

Except that there was no significant difference a day after spraying treatments from the same scion-stock combination, the contents of ABA and ZT in buds and leaves of A1, A2 and A3 from 16 and 31 days after spraying were significantly higher than those of CK1, CK2 and CK3, whereas B1, B2 and B3 were significantly lower, or no significant difference, respectively (Fig. 5A-D). The contents of IAA also were non-significant a day after spraying treatments from the scion-stock combination. Furthermore, the contents of IAA in buds and leaves of B1, B2 and B3 from 16 and 31 days after spraying were significantly higher than those of CK1, CK2 and CK3, whereas A1, A2 and A3 were significantly lower, respectively (Fig. 5E, F). Moreover, the contents of ABA, ZT and IAA in buds and leaves of all treatments increased from day 1 to 31 days. Besides, the overall trend of endogenous hormone contents in buds and leaves of the three scion-stock combinations under the same treatment also was dwarf self-rooted rootstock M.26, followed by the interstock M.26/*M. sieversii*, and finally the vigorous rootstock *M. sieversii*.

**Fig. 5.**
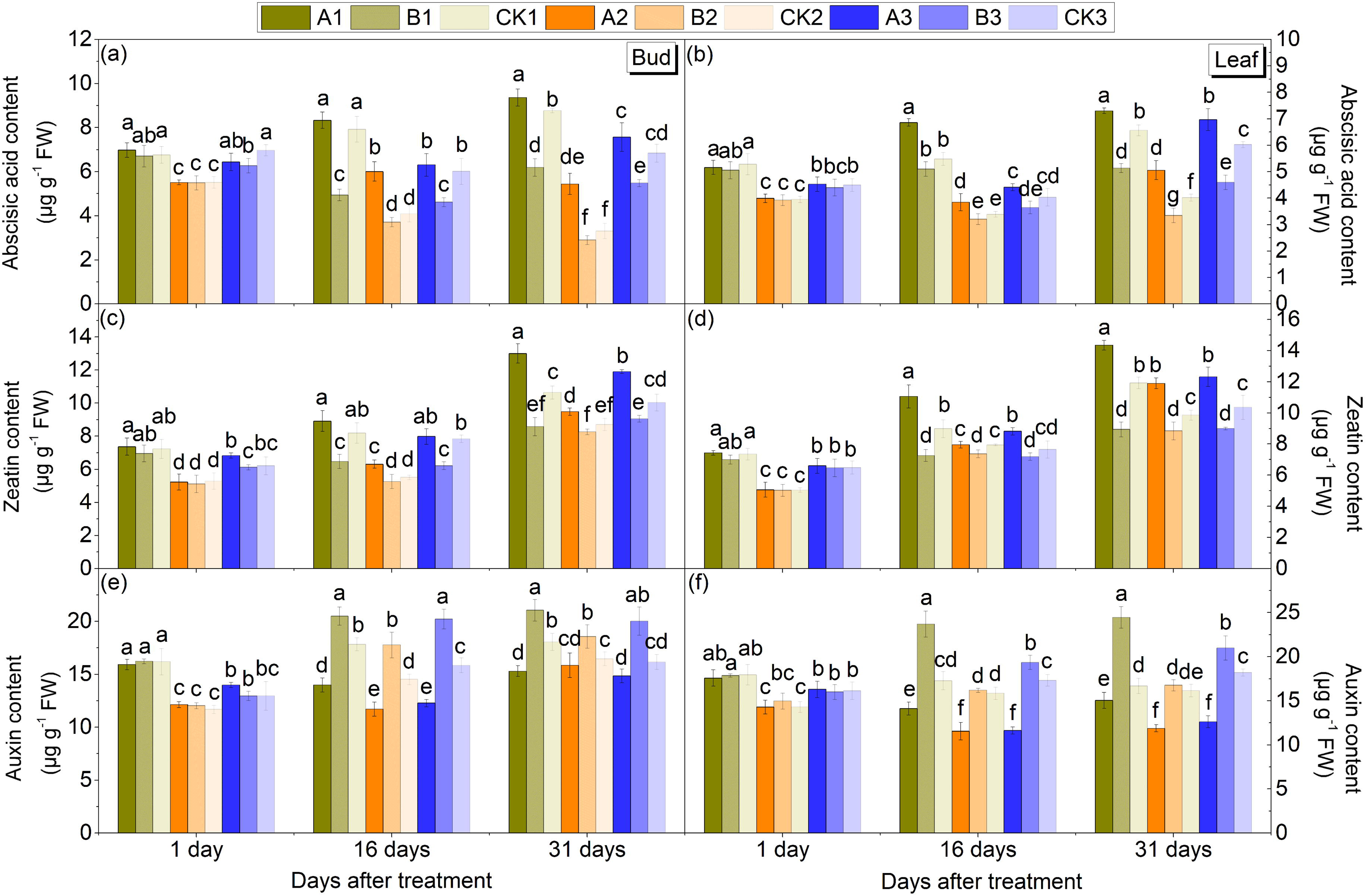
Changes of endogenous ABA (a and b), ZT (c and d) and IAA (e and f)contents in buds and leaves of 3 scion-stock combinations including Nagafu No.2/M.26, Nagafu No.2/*M. sieversii* and Nagafu No.2/M.26/*M. sieversii* after exposure to 800 mg·L^−1^ 2-Chloroethylphosphonic acid (CEPA) and 2 μL·L^−1^ its inhibitor of 1-methylcyclopropene (1-MCP) after 1, 16 and 31 days. Bars indicate SE. Different small letters within the figures indicate significant difference at 0.05 levels.

### Effects of CEPA and 1-MCP on the flowering genes expression level in buds and leaves of apple from scion-stock combinations

After 16 days of spraying with CEPA, the expression level of the floral negative regulator gene *TFL1* was significantly down-regulated, and the floral positive regulator gene *AP1* was significantly up-regulated in buds of dwarf self-rooted rootstock M.26 and the interstock M.26/*M. sieversii* compared with the control, whereas the effect on the vigorous rootstock *M. sieversii* was not significant (Fig. 6). Furthermore, there was no significant difference in leaves. However, the expression levels of *TFL1* and *AP1* were not significantly different from the control after the application of 1-MCP.

**Fig. 6.**
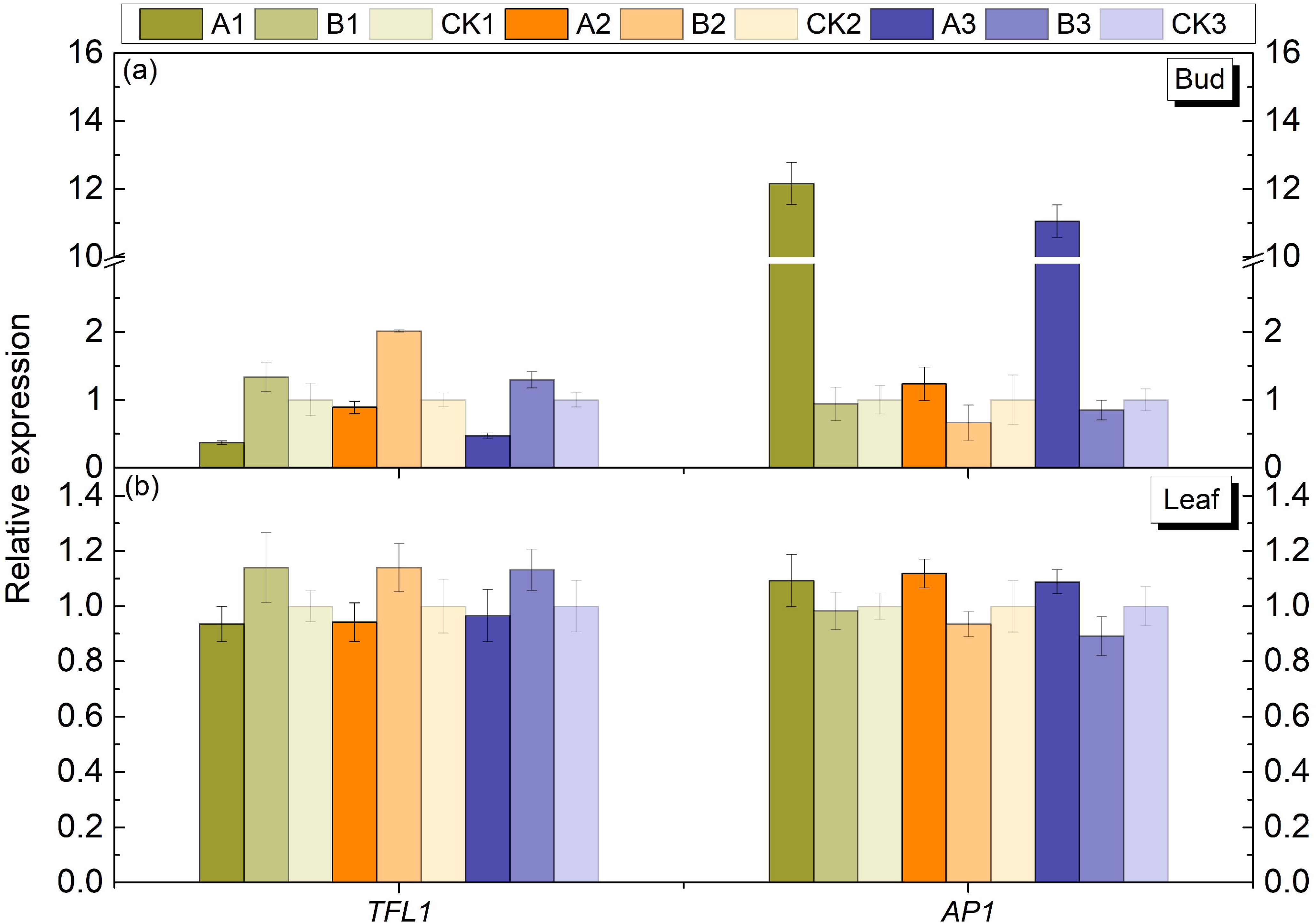
Changes of the relative expression levels of 2 flowering-related genes *TFL1* and *AP1* from leaves (a) and buds (b) of 3 scion-stock combinations including Nagafu No.2/M.26, Nagafu No.2/*M. sieversii* and Nagafu No.2/M.26/*M. sieversii* after exposure to 800 mg·L^−1^ 2-Chloroethylphosphonic acid (CEPA) and 2 μL·L^−1^ its inhibitor of 1-methylcyclopropene (1-MCP) after 16 days. The left y-axis indicates relative gene expression levels were determined by qRT-PCR and analyzed using 2^−ΔΔC^T Method. All qRT-PCR for each gene used three biological replicates, with three technical replicates per experiments; error bars indicate SE. Different lower case letter indicates the significant difference among four treatments at *P* = 0.05.

## Discussion

### CEPA promoted flower induction mainly by promoting the accumulation of carbohydrates, ABA and ZT

Ramirez and Hoad (1981) suggested that CEPA could inhibit growth and promote flower, probably due to CEPA inhibited the biosynthesis and operation of IAA and GA in shoots. The result of flower bud morphology in this study showed that flower bud differentiation of apple cv. ‘Nagafu No.2’ was promoted at 16 days and 31 days after spraying with CEPA compared with the control, while the effect of 1-MCP on flower bud differentiation was not significant (Fig. 1A, B). The flowering rate also was significantly increased and the blossom period was 2 days ahead of time compared with the control after spraying with CEPA, whereas the flowering rate was significantly decreased and the blossom period was delayed for 2 days after spraying with 1-MCP. Accordingly, the content of carbohydrates from buds have a significant positive impact on the flower bud formation in fruit trees (Garcia-Lui *et al*., 1995; Shalom *et al*., 2014). Wünsche *et al*. (2005) showed that the content of various monosaccharides and starch in leaves and terminal buds from 7-year-old apple cv. “Braeburn/M26” changed significantly with the passage of time. Xing *et al*. (2014) found that the accumulation of sucrose, glucose, fructose and soluble sugars during flower induction can promote flower bud differentiation as an energy substance in buds and leaves of “Fuji” apple. In the current work, the starch contents in buds and leaves from A1, A2 and A3 were significantly decreased at 16 days and 31 days after spraying with CEPA, whereas slightly increased from B1, B2 and B3 after the 1-MCP application in contrast to those of CK1, CK2 and CK3, respectively (Fig. 3A, B). Otherwise, the contents of soluble sugar, glucose and sucrose were significantly promoted at 16 days and 31 days after spraying with CEPA, and slightly inhibited after the 1-MCP application (Fig. 3C-H). Suggesting that CEPA can promote the synthesis and accumulation of soluble sugar, glucose and sucrose in buds and leaves of apple cv. ‘Nagafu No.2’, but inhibit the synthesis and accumulation of starch, therefore, promote flower bud induction as an energy substance. Meanwhile, the activities of α-AMY and SPS also were increased at 16 days and 31 days after spraying with CEPA, and slightly decreased after the 1-MCP application, which were contrary to the activities of SS in buds and leaves of apple cv. ‘Nagafu No.2’ (Fig. 4A-F). These results were consistent with previous studies that the starch content decrease was accompanied by a small increase in the a-AMY activity (Lambrechts *et al*., 1994), an SPS activity negatively related with starch content in leaves but did not inhibit normal plant growth (Hashida *et al*., 2016), and there was a highly significant negative correlation observed between the SS activity and sucrose content in the florets and branchlets of broccoli (*Brassica oleracea* L.) (Pramanik *et al*., 2004).

Researchers have done a lot of research on the regulation of GA on flower bud differentiation in fruit trees (Xing *et al*., 2014; Yamaguchi *et al*., 2014). Apart from GA, several other hormones such as ABA, auxin, CTK and ETH are closely related to flower bud differentiation (Bangerth, 2008; Grochowska and Hodun, 2012). In the present research, the increment of terminal shoot length and new shoot length was significantly inhibited after spraying with CEPA, whereas slightly promoted or no significant difference after spraying with 1-MCP (Table 1). Simultaneously, the contents of ABA and ZT were increased, while IAA was decreased after spraying with CEPA, which contrary to 1-MCP (Fig. 5A-F). This is mainly because 1-MCP strongly competes with ETH receptors after spraying 1-MCP and binds to ETH receptors through metal atoms, thus blocking the normal binding of ETH to its receptors. Consequently, we concluded that the application of exogenous CEPA promoted flower bud differentiation by promoting the synthesis and accumulation of ABA and ZT and inhibiting the accumulation of IAA, thereby suppressing vegetative growth and promoting reproductive growth.

### CEPA was more susceptible to easy-flowering rootstock and still less susceptible to difficult-flowering rootstock

Growth of the scion was affected by the genotype of the rootstock (Dieleman *et al*., 1998; Gonçalves *et al*., 2006). According to Richards *et al*. (1986), the content of GA in leaves and branches from interstock of the dwarfing cultivar M9 decreased compared to the nondwarfing M115 interstock. Kamboj *et al*. (1999) showed that the contents of CTK in the branches and roots from the vigorous rootstock MM106 were higher than those from the dwarfing rootstocks, M26 and M9. Similarly, the overall trend of the increment of terminal shoot length, new shoot length and new shoot diameter from the three different scion-stock combinations under the same treatment in this study was the vigorous rootstock *M. sieversii*, followed by the interstock M.26/*M. sieversii*, and finally the dwarf self-rooted rootstock M.26. It was consistent with IAA content while contrary to the content of ABA and ZT (Fig. 5), thus conducive to vegetative growth and the flowering rate showed the opposite trend (Fig. 2). The highest rate of flowering was from dwarf self-rooted rootstock M.26, followed by interstock M.26/*M. sieversii*, and finally the difficult-flowering rootstock, vigorous rootstock *M. sieversii*. Furthermore, after the application of CEPA and 1-MCP, the flowering rates of A1 and A3 were significantly increased in contrast to those of CK1 and CK3, while those of B1 and B3 were significantly decreased, but no significant difference was found between A2, B2 and CK2.

Collectively, the application of CEPA and 1-MCP was more susceptible to easy-flowering rootstock dwarf self-rooted rootstock M.26, followed by interstock M.26/*M. sieversii*, and still less susceptible to difficult-flowering rootstock, vigorous rootstock *M. sieversii*.

### Flower bud induction is closely linked to flowering-related genes TFL1 and AP1 in buds and leaves of apple from scion-stock combinations

Liljegren *et al*. (1999) reported that the normally sharp phase transition between the production of leaves with associated shoots and formation of the flowers in Arabidopsis, which occurs upon floral induction, is promoted by positive feedback interactions between *LEAFY* (*LFY*) and *APETALA1* (*AP1*), together with negative interactions of these two genes with *TERMINAL FLOWER1* (*TFL1*). *TFL1* is a key negatively regulator of flowering time and the development of the inflorescence meristem in *Arabidopsis thaliana* (Hanano and Goto, 2011). In this study, the expression of *TFL1* was repressed while *AP1* was promoted in buds and leaves in contrast to the control after 16 days of spraying with CEPA (Fig. 6). However, the application of 1-MCP has the opposite effect. In concluded, the application of CEPA can up-regulate the expression of genes that are flowering positive regulators and down-regulate the expression of genes that are flowering negative regulators, resulting in the promotion of flower formation. In addition, there was no significant difference or significant inverse effect between 1-MCP and control compared with CEPA. This results are consistent with the findings that treatment with 1-MCP and ETH generally produced opposite effects on related genes (Yang *et al*., 2016).

## Acknowledgments

This research was financially supported by the Fostering Foundation for the Excellent Ph.D. Dissertation of Gansu Agricultural University (2017002), and the Key Scientific Technology Research Projects of Gansu Province (GPCK2013-2).

